# Fezf2 regulates differentiation of Aire-expressing and post-Aire mimetic epithelial populations maintaining thymic homeostasis

**DOI:** 10.64898/2025.12.31.697099

**Authors:** Joe Germino, Xian Liu, Yi Wang, Jared Balolong, Vera Tang, Im-Hong Sun, Yanzhen Nie, Jeremiah Tsyporin, Irina Proekt, Kristin Rattay, Vasilis Ntranos, Mark S. Anderson, Bin Chen, James M. Gardner, Corey N. Miller

## Abstract

Fezf2 has been proposed to serve principally as a transcriptional regulator of broad self-antigen expression in medullary thymic epithelial cells (mTECs). Here, we demonstrate an additional function for Fezf2 as a developmental regulator of TEC lineage commitment. Fezf2 deficiency in the thymus led to a relative expansion of lung mimetic and Ccl21+ immature mTECs at the expense of MHCII-hi Aire-expressing and all other mimetic populations. Consistent with this, the disruption of mTEC development had functional consequences, including alterations in tuft-associated iNKT polarization and microfold-associated B-cell class switching. In addition, high-resolution transcriptomics revealed that Fezf2 and Aire regulate distinct transcriptional programs, with Fezf2 driving both activation and repression of a more limited set of tissue-restricted genes compared to the broad gene activation program observed for Aire. Pure transcriptional repression at Fezf2 target loci was sufficient to partially rescue mTEC development in global Fezf2-/- (tKO) mice, potentially through suppression of Lifr expression and modulation of downstream Stat3 signaling tone. These results expand our understanding of Fezf2 in mTEC biology, highlighting transcriptional repression as a required functional facet for adult steady-state lineage patterning across the mTEC compartment.

## INTRODUCTION

The thymic medulla plays an essential role in T-cell development by enforcing stringent negative selection of developing single-positive thymocytes. This requires broad proteomic expression, abundant antigen processing, and antigen display on major histocompatibility proteins to screen for potential self-reactivity of individual, somatically recombined TCRs. To facilitate this, medullary thymic epithelial cells (mTECs) have evolved to express near-complete coverage of the self-proteome. Two complementary mechanisms are understood to enable proteomic coverage across mTECs: direct semi-stochastic genome-wide promiscuous gene expression (PGE), as has been suggested for the autoimmune regulator protein (Aire), and biologically ordered cellular differentiation of diverse epithelial states, as for lineage defining transcription factors (1–5). The balance between these mechanisms and their impact on broader thymic function remain active areas of investigation, but recent advances in “omics” sequencing technologies have enabled rapid advances in our understanding of both.

In addition to Aire, the zinc-finger transcription factor Fezf2 has been proposed to directly induce expression of a unique set of tissue restricted antigens (TRAs) within mTECs (6, 7). However, the original descriptions of altered TRA expression in Fezf2-/-mice relied on bulk microarray analysis of mTEC populations that have since been revealed to contain heterogeneous subsets of terminally-differentiated transcriptional states (8–10). Recently, it was reported that loss of Fezf2 in mTECs alters thymic tuft cell development, suggesting that Fezf2 has roles in mTEC biology beyond induction of cell-intrinsic direct transcriptional activation for TRA expression (11, 16). Outside the thymus, Fezf2 is required for cell-fate determination during neuronal development, where it acts as a transcriptional repressor, recruiting Tle corepressors through its Eh1 motif (12). This repressive activity is crucial for restricting corticothalamic and callosal cortical neuron gene programs, enabling normal subcerebral neuron development (13–15). The degree to which Fezf2 may similarly function as a transcriptional repressor of lineage commitment in mTEC biology has not been extensively explored (6, 7, 11, 16).

Here we leveraged single-cell genomics and a range of functional approaches to further define the role of Fezf2 in mTEC biology. We find that Fezf2 alters steady-state mTEC developmental dynamics, with a relative expansion of Ccl21+ immature and lung “mimetic” cells at the expense of thymic tuft cells and Aire-expressing lineages, including most other post-Aire terminally differentiated mimetic subsets (9, 11). Transcriptionally, the Fezf2 program is distinct from that of Aire, and is characterized by a more restricted program of both gene activation and repression (7, 16). Indeed, pure transcriptional repression directed to Fezf2 target loci was sufficient to partially rescue mTEC developmental dynamics in Fezf2-deficient mice. Mechanistically, Lifr was amongst the most Fezf2-repressed gene targets and was prominently expressed in Ccl21+ and lung mimetic subsets in Fezf2 cKO mice, suggesting a possible role for Stat3 signaling tone in establishing the Fezf2 cKO phenotype. Overall, this work underscores the essential role of Fezf2 in shaping mTEC composition and maintaining the functional integrity of the thymic microenvironment.

## RESULTS

### Fezf2 deficiency results in broad alterations in steady-state mTEC development

To better understand the role of Fezf2 in mTEC biology, we sorted two independent samples of Epcam+CD45-TECs each from five pooled 4- to 6-week-old FoxN1-Cre+/-; Fezf2fl/fl (Fezf2 cKO) or Aire-/- (Aire KO) C57BL/6 thymi along with age and sex-matched pooled Foxn1-Cre-/-; Fezf2fl/fl and Aire+/+ (WT) control thymi for 10x Chromium scRNA-seq (**Fig. S1A, 1A**). Because high resolution alterations in mTEC development and heterogeneity have been described for Aire KO mice, we also included Aire KO samples in our dataset as a control and comparator (9). These datasets were of high quality with a total of 22,993 WT, 12,951 Fezf2 cKO, and 14,448 Aire KO cells remaining after processing and filtering steps (**Fig. S1B-C**). Normalization and batch correction with scvi-tools produced well-integrated samples for downstream analysis (**Fig. S1D**)

(17). To standardize nomenclature with published datasets, we utilized celltypist to transfer published high-resolution labels to our dataset (**Fig. S1E, 1B**) (9, 18). Using this approach, we identified nearly all adult mTEC subsets in our dataset, including ‘Adult cTEC’, ‘Transit-amplifying (TA) mTEC’, ‘Immature’, ‘Aire-stage’, and multiple mimetic subsets, but not Myog-expressing muscle or Ptf1a-expressing pancreatic mimetic cells (**Fig. S1E**). Observation of these two minor mimetic subsets likely requires enrichment for post-Aire mTECs, which was not performed here. After label transfer, we noticed that the immature mTEC subset in our dataset appeared to contain multiple transcriptionally distinct subpopulations, so we performed subclustering analysis to further resolve this heterogeneity into three subpopulations (**Fig. S1F, 1B**). To identify unique markers associated with these subsets, we performed differential gene expression analysis and found that they were easily distinguished by expression of Ccl21a, Bcam, and Xylt1, respectively (**Fig. S1F, Table S1**). Bcam was recently described to mark a subset of epithelial progenitors in the human thymus, Xylt1 is a xylosyltransferase required for proteoglycan biosynthesis and collagen extracellular matrix remodeling, and Ccl21a is a described marker of immature mTECs in other studies (19, 20). Next, we examined expression of Aire and Fezf2 across our dataset. Consistent with previous work, Aire expression was detected at low levels in cycling transit-amplifying (TA) mTECs before peaking in Aire-stage mTECs, with no Aire expression detected in mimetic cell populations or immature mTECs (**Fig. 1C**) (21). In contrast, Fezf2 expression was observed in two waves, the first occurring during the transition between TA mTECs and Aire-stage mTECs and the second during late Aire expression and the transition to post-Aire mimetic lineages (**Fig. 1C**). Notably, Fezf2 expression extended into some Ccl21a-expressing immature mTECs and into a subset of cells annotated as basal skin and basal lung mimetic cells, suggesting roles for Fezf2 expression before and after Aire expression (**Fig. 1C**). The chromatin remodeler Chd4, which has been implicated in co-regulating Fezf2- and Aire-induced TRA expression, was broadly expressed at modest levels across mTECs in our dataset (**Fig. S1G**) (7).

**Figure 1.**
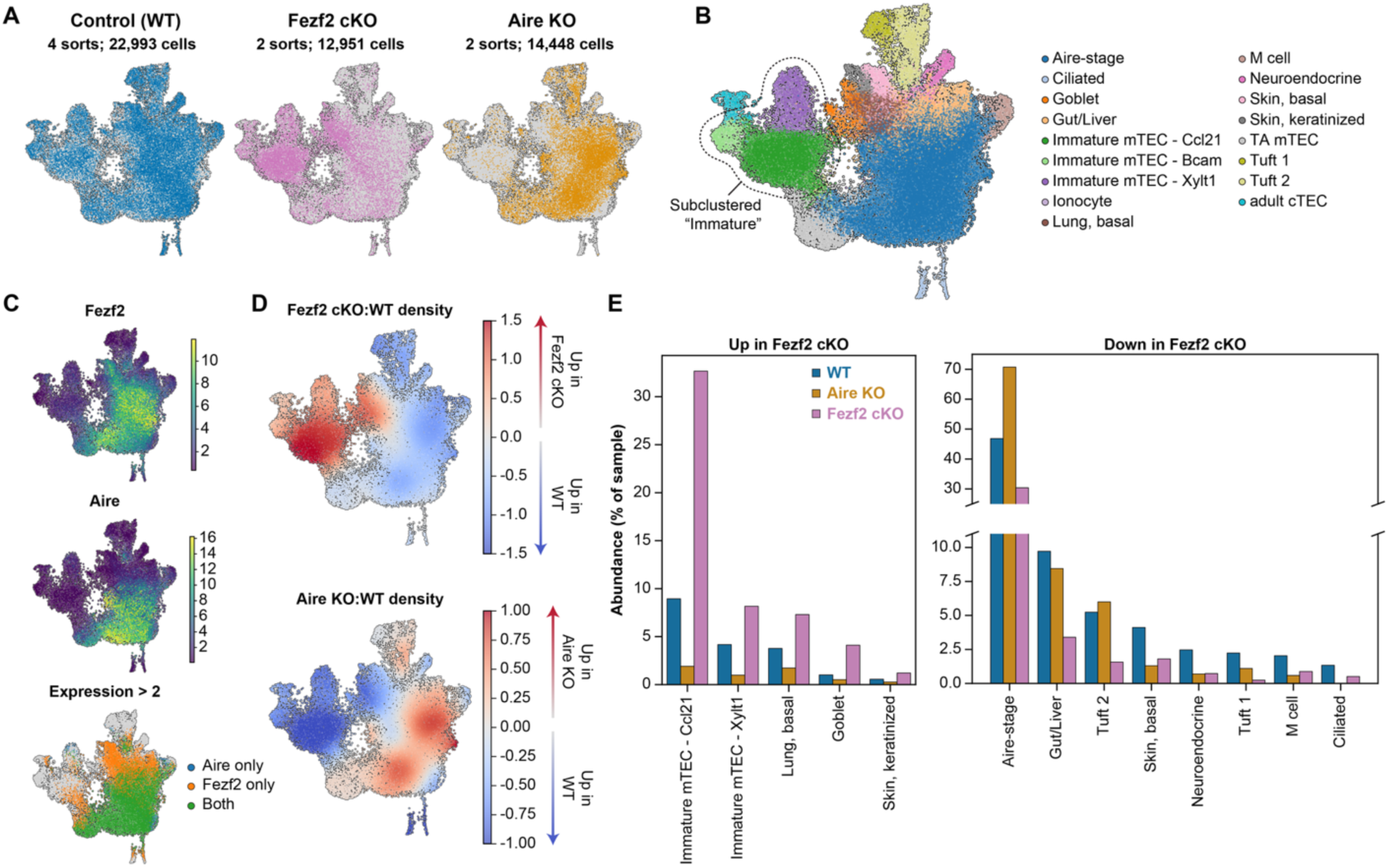
Fezf2 deficiency results in broad alterations to steady state mTEC development. By scRNA-seq. (**A.**) UMAP depicting cells colored by genotype of scRNA-seq samples collected from 4–6-week-old WT, Aire KO, and Fezf2 cKO mice. (**B.**) UMAP depicting cells colored by cell type of scRNA-seq samples. (**C.**) Feature plots depicting normalized expression (top, middle) and thresholded normalized expression > 2 (bottom) of Aire and Fezf2 in WT TECs. (**D.**) Differential density plot showing increasing (red) or decreasing (blue) populations in the Aire KO or Fezf2 cKO samples relative to WT. (**E.**) Fraction of cells annotated as each TEC subpopulation across the WT, Aire KO, and Fezf2 cKO samples for subpopulations with a mean log2(fold change) between WT and Fezf2 cKO samples greater than 0.5 or less than -0.5. Colored bar depicts the mean abundances across 2 (Fezf2 cKO and Aire KO) or 4 (WT) independent replicates.

Next, we wished to understand how loss of Fezf2 alters the adult steady-state distribution of mTEC transcriptional states. Differential density analysis of the relative abundance of TEC populations in Fezf2 cKO mice revealed a pronounced relative expansion of immature mTECs as well as goblet and basal lung mimetic cells, but a decrease across most other subsets (**Fig. 1D**). Reassuringly, differential density analysis of Aire KO mice produced the expected and previously reported loss of most mimetic subsets with a corresponding expansion of putative Aire-stage mTECs, providing orthogonal validation of the technical coherence of our datasets (**Fig. 1D**) (9). As a percent of all TECs in each sample, Ccl21a-expressing immature mTECs increased from approximately 10% of WT samples to over 30% of Fezf2 cKO samples while the tuft cell subsets decreased from 2-5% of WT to 1.5% or less of Fezf2 cKO samples, consistent with recent reports describing loss of thymic tuft cells in Fezf2-deficient mice (**Fig. 1E**) (11, 16). The frequencies of Aire-stage and other mimetic cells were also considerably reduced in Fezf2 cKO samples but were less severely impacted, and transit-amplifying mTECs were present at a similar frequency across genotypes (**Fig. 1E, S1H**). Critically, we validated this pattern of disrupted adult steady-state mTEC development in Fezf2 cKO mice by flow cytometry using a panel of standard mTEC and described mimetic cell markers (**Fig. 2A-B, Table S2**). In contrast to previously published data, but consistently across our study, Ccl21-expressing mTECs were substantially increased in frequency and MHCII-hi Aire+ mTECs decreased in frequency in Fezf2 cKO thymi (6, 16). The absolute number of Ccl21+ cells was not statistically different between genotypes, as expected given that the total number of mTECs in cKO thymus was decreased (**Fig. 2A-B**). Collectively, these observations reveal broad compositional shifts across the mTEC compartment in Fezf2 cKO mice and expand the role of Fezf2 in steady-state adult mTEC development.

**Figure 2.**
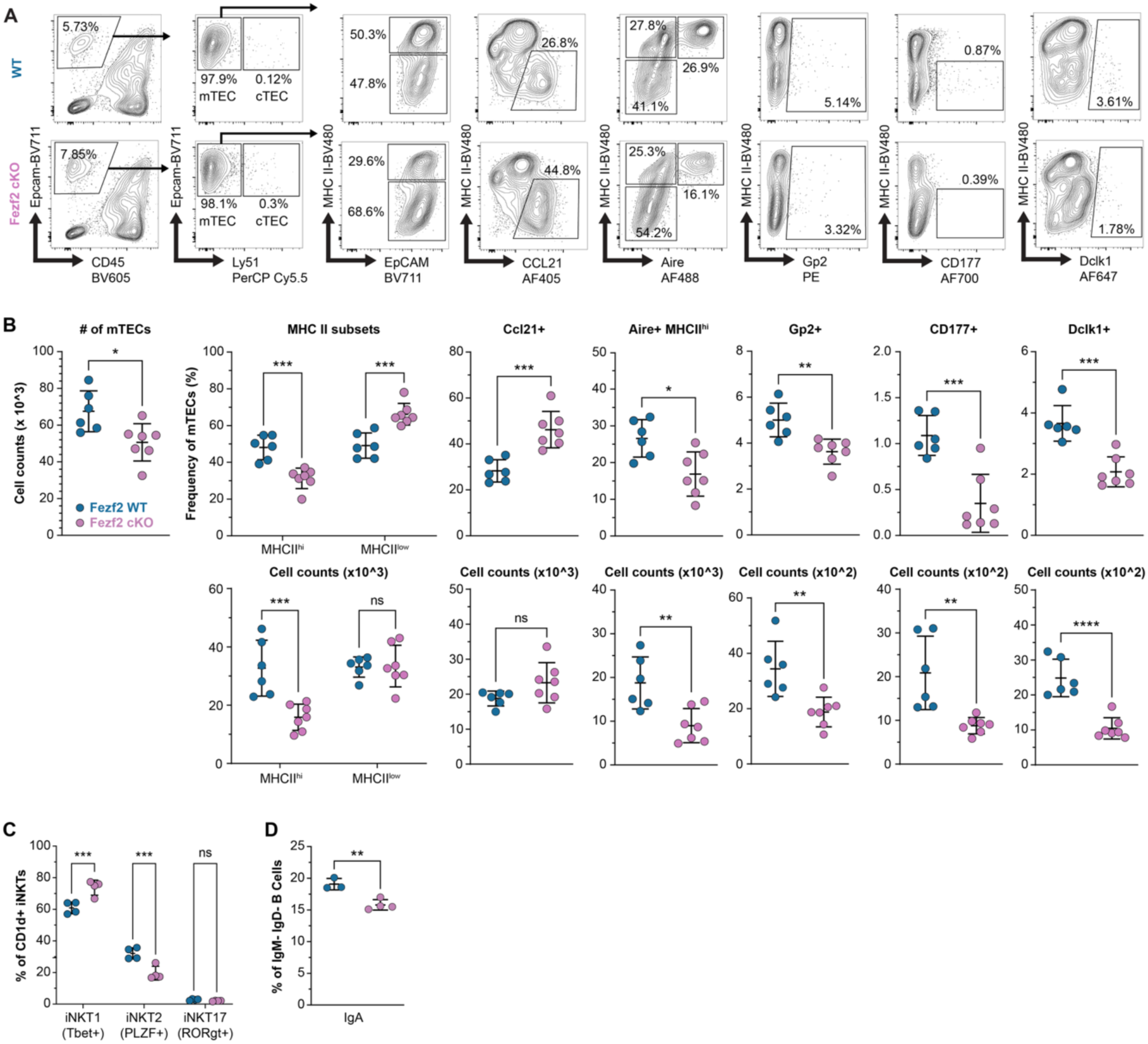
Fezf2 deficiency results in broad alterations to steady state mTEC development by flow cytometry. (**A.**) Representative gating strategy for identifying major mTEC subpopulations by flow cytometry in 6-week-old WT and Fezf2 cKO mice. (**B.**) Frequency and number of total mTECs and major mTEC subpopulations across 6-week-old WT (n=6) and Fezf2 cKO mice (n=7), determined by flow cytometry. (**C.**) Frequency of thymic iNKT cell subsets in 6-week-old WT (n=4) and Fezf2 cKO (n=4) mice. (**D.**) Frequency of thymic IgM- IgD- IgA+ B cells in 6-week-old WT (n=4) and Fezf2 cKO (n=4) mice. Statistical significance was calculated using unpaired Student’s t test (B-D).

### Developmental defects in Fezf2 cKO thymus disrupt stromal-immune crosstalk

Because the observed alterations to mTEC development in Fezf2 cKO mice were more extensive than expected, we next sought an orthogonal means of interrogating epithelial function. Thymic tuft and M cells have been shown to contribute to normal development of iNKT cells and thymic B cell maturation, respectively (9, 10, 22). Therefore, we asked whether these mimetic functions were disrupted in cKO thymus. Indeed, the number and frequency of iNKT2 cells was considerably decreased in Fezf2 cKO mice relative to WT, as would be expected with a pronounced defect in thymic tuft cell development (**Fig. S2A, 2C**) (22, 23). Notably, there was a concomitant increase in the frequency, but not number, of Tbet+ iNKT1 cells, which is consistent with an expanded Ccl21+ compartment and increased IL-15 trans-presentation (**Fig. S2A, 2C**) (23, 24). Next, we turned to the thymic B cell compartment, which has been shown to depend on M cell-derived Ccl20 for medullary organization, maturation, and IgA class-switching (10). Accordingly, in Fezf2 cKO thymus, IgA class switching was diminished by flow cytometry (**Fig. S2B, 2D**). To investigate this further, we examined the physical association between B220+ cells and Gp2+ M cells. In WT tissue, most Gp2+ cells were directly touching B220+ cells or were within 20um of a B220+ cell surface (mean distance = 7.02um, 93.3% within 20um) by quantitative image analysis (**Fig. S2C**). However, this distance increased significantly (mean = 18.08um) for Gp2+ cells in Fezf2 cKO thymus, with only 67.5% of Gp2+ cells within 20um of the nearest B cell (**Fig. S2C**). In combination, these findings align with disrupted mTEC development in Fezf2 cKO mice and reveal altered medullary function secondary to loss of specific mimetic populations.

### Fezf2 and Aire regulate distinct transcriptional programs

To better understand the transcriptional program mediated by Fezf2 in mTECs, we first compared it to the program regulated by Aire. We identified differentially expressed (DE) genes across genotypes within each mTEC subpopulation (**Table S3-4**). Among the Fezf2 DE genes in our dataset were many genes previously identified as Fezf2-regulated using bulk microarray data, indicating our results are consistent with previous studies (**Fig. 3A**) (6). A majority of both Aire and Fezf2 DE genes were classified as TRAs in our analysis, reinforcing that both transcription factors regulate genes associated with specialized cell states (**Fig. S3A-D, 3B**). Consistent with previous reports, Aire overwhelmingly induced gene expression in our dataset, while Fezf2 regulated a more balanced program that included a substantial number of repressed genes (**Fig. 3C**) (7, 16). Utilizing our Fezf2 cKO and Aire KO single-cell datasets, we next quantified the number of genes expressed per cell as a metric for the transcriptional diversity of individual cells. We examined changes in transcriptomic coverage of mTEC subpopulations across genotypes and found that while Aire KO Aire-stage mTECs had a profound decrease in the number of genes expressed per cell, Fezf2 cKO Aire-stage mTECs had only a modest decrease (**Fig. 3D**). We then quantified the fraction of these expressed genes that are TRAs per cell based on Tau and Shannon entropy scores calculated for each gene from its distribution of expression across cell types in the cellxgene single-cell mouse tissue atlas (**Fig. S3A-D and Table S5**). As expected, Aire-stage mTECs had a marked decrease in the fraction of expressed genes annotated as TRAs in the Aire KO thymus; by contrast, in the Fezf2 cKO thymus the fraction of TRAs among expressed genes was only subtly affected (**Fig. 3D**). The loss of transcriptomic coverage in Aire-deficient but not Fezf2-deficient Aire-stage mTECs was also apparent in the magnitude of gene expression across genotypes, with a substantial decrease in the average log-transformed counts of many lowly-expressed genes in Aire KO, but not Fezf2 cKO, Aire-stage mTECs compared to WT (**Fig. 3E**).

**Figure 3.**
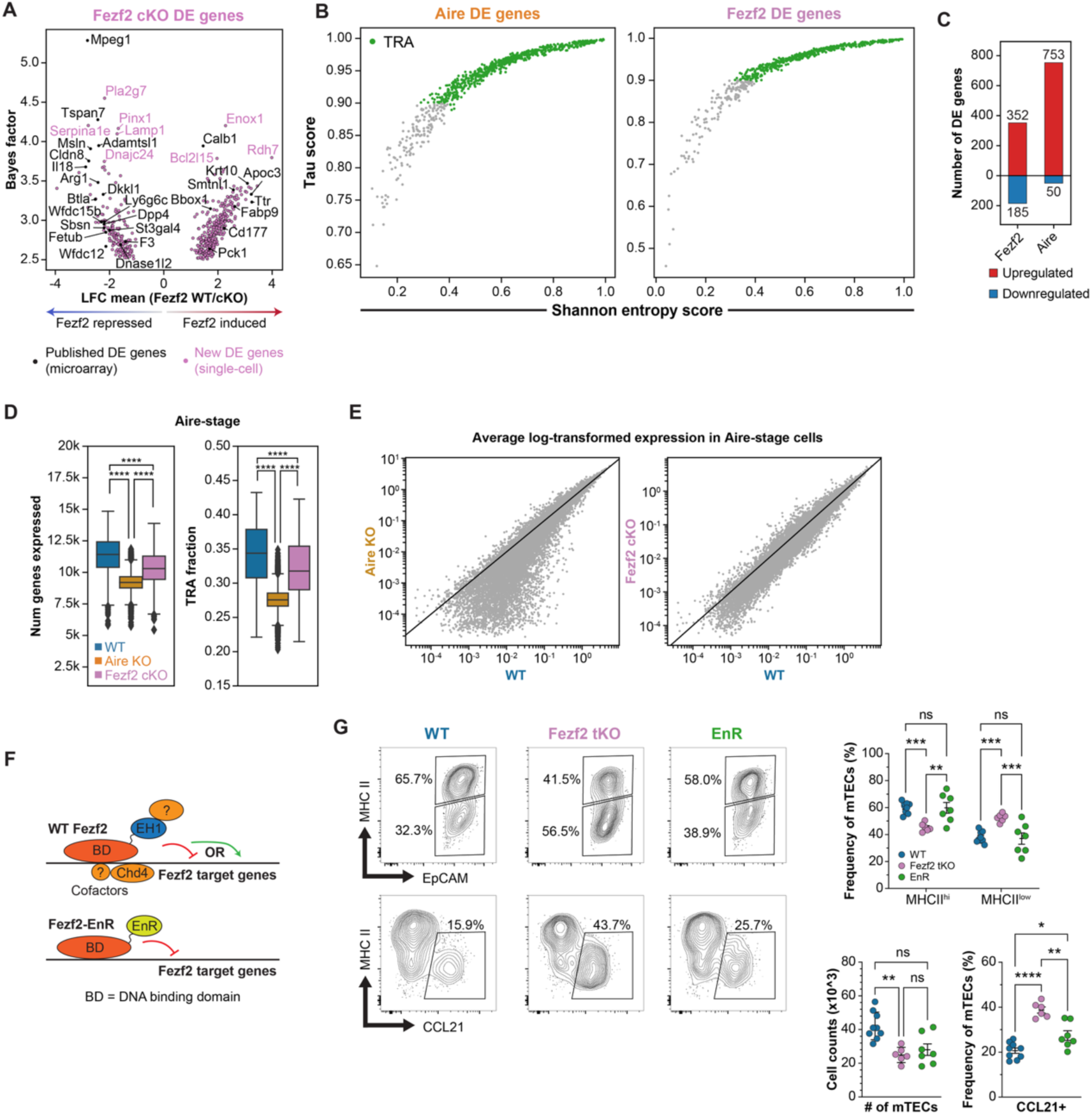
Aire, Fezf2 regulate distinct programs of transcriptional diversity. (**A.**) Mean log fold change and bayes factors for Fezf2-regulated genes that were identified as highly differentially expressed between cKO and WT samples in each mTEC subset (**see methods**). (**B.**) Shannon entropy vs Tau scores of Fezf2 and Aire DE genes, highlighting genes annotated as TRAs. (**C.**) Number of Aire or Fezf2 differentially expressed genes that are upregulated (red) or downregulated (blue) by their respective transcription factor. (**D.**) Number of genes with scVI normalized expression greater than 0.1 in Aire-stage mTECs across genotypes (left) and fraction of expressed genes annotated as TRAs (right). (**E.**) Average log-transformed expression of all genes in Aire-stage mTECs between WT, Aire KO, and Fezf2 cKO samples. Diagonal line marks equivalent expression in WT and KO samples. (**F.**) Schematic depicting effect of transgenic Fezf2-EnR protein on Fezf2 target genes compared to WT Fezf2. (**G.**) Representative flow cytometry gating (left) and frequency of major mTEC subpopulations across 4- to 7-week-old WT, Fezf2 tKO and Fezf2 tKO EnR mice (right). (n=6-9 mice). Statistical significance was calculated using two-way ANOVA (Sidak’s test) with multiple comparisons (G).

Fezf2 has a well-established role as a direct transcriptional repressor in neuronal development, where it recruits the co-repressor Tle4 to its target genes to modulate cell fate decisions (12, 25, 26). Therefore, we hypothesized that Fezf2-mediated transcriptional repression might also regulate mTEC differentiation programs, thereby contributing to TRA coverage through access to terminally differentiated epithelial states. Previous work on the role of Fezf2 in neuronal cell-fate decisions has demonstrated that pure transcriptional repression at Fezf2 loci substantially rescues the neurodevelopmental defects in mice lacking Fezf2 (**Fig. 3F**) (12). This experimental system utilized transgenic expression of the Drosophila engrailed repressor domain fused to the Fezf2 DNA binding domain, expressed under the control of the Fezf2 promoter (Fezf2-EnR transgenic allele). Crossing the EnR allele to Fezf2-/- (Fezf2 tKO) mice allowed evaluation of the effect of isolated direct repressive function at Fezf2 target loci on brain development. Therefore, we leveraged this system to investigate if direct transcriptional repression is also sufficient to rescue adult steady-state mTEC developmental dynamics in the absence of Fezf2. First, flow cytometry confirmed that Fezf2 tKO mice had a similar pattern of disrupted mTEC development as was observed in Fezf2 cKO mice (**Fig. S3E-G, 3G**). Next, we compared mTEC subsets in WT mice to those in Fezf2 tKO EnR mice by flow cytometry. Strikingly, we observed a near-complete normalization of both the frequency and number of Ccl21- and MHCII-expressing subsets in Fezf2 tKO mice carrying the repressive Fezf2-EnR transgenic allele (**Fig. S3F, 3G**). Interestingly, Fezf2 tKO EnR mice did not exhibit a rescue of normal mimetic cell subcompartments, raising the possibility that at least some mimetic developmental pathways require direct transcriptional induction by Fezf2 and its thymic binding partners (**Fig. S3G**). In total, these data suggest that Fezf2 is not primarily acting alongside Aire in MHCII-hi mTECs to broadly induce TRA gene expression but that it has a prominent role in mTEC differentiation that relies on direct transcriptional repression.

### Lifr is a Fezf2-repressed target gene

To better understand how the Fezf2 transcriptional program could be affecting mTEC development we looked for genes with known roles in cellular plasticity and differentiation across the Fezf2-regulated genes identified in our scRNA-seq dataset. Among the top Fezf2 transcriptionally repressed genes was Lifr, an IL-6 family cytokine receptor that forms a functional complex with Gp130 (Il6st) to initiate downstream Stat3 signaling often associated with cellular plasticity or stem cell renewal processes (**Fig. 4A**) (27–29). When mapped across mTEC subsets in our scRNA-seq dataset, Lifr was found to be highly expressed in both WT and Fezf2 cKO immature mTECs but was dramatically upregulated in a subset of lung mimetic epithelial cells in Fezf2 cKO mTECs (**Fig. 4B-C**). Notably, the mTEC populations with high Lifr expression were also those subsets that were overrepresented in Fezf2 cKO thymi by scRNA-seq (**Fig. 1D-E**). To validate Lifr expression by flow cytometry, we identified differentially expressed marker genes for the putative goblet-like or basal lung-like cells vs all other cKO mTECs in our scRNA-seq dataset (**Fig. 4D**). Surprisingly, among these marker genes was Gp2, which has been used as a specific marker for thymic M cells but is also known to be expressed by respiratory epithelial cells (9-10, 30) (**Fig. S4A**). Krt6a was also largely restricted in its expression to both goblet and basal lung mimetic cells in our Fezf2 cKO scRNA-seq samples (**Fig. S4A**). In combination, Krt6a was found to be expressed by a subset of Gp2+ cells and both the frequency and number of Gp2+Krt6a+ mTECs was increased in Fezf2 cKO thymi, confirming that Krt6a-expressing mTECs annotated as goblet and basal lung-like epithelial cells are overrepresented in Fezf2 cKO mice (**Fig. S4B-C**). Interestingly, most Gp2-expressing cells with close association to Krt6 were spherical in shape and morphologically distinct from the canonical clefted structure of WT thymic M cells (**Fig. S4D**). By flow cytometry, both the fraction and absolute number of total Lifr+ mTECs and Lifr+ Ccl21-Gp2+ lung mimetic cells were increased in Fezf2 cKO relative to WT mice (**Fig. S4E, 4E**). Finally, to investigate possible sources of Lif cytokine in the murine thymus we reanalyzed a publicly available scRNA-seq dataset containing the major intrathymic hematopoietic cell populations before and after dexamethasone treatment (**Fig. S4F**) (31). In this dataset, Lif transcript was readily detected within thymic ILCs, early thymic progenitors (ETPs), and gamma delta T cells.

**Figure 4.**
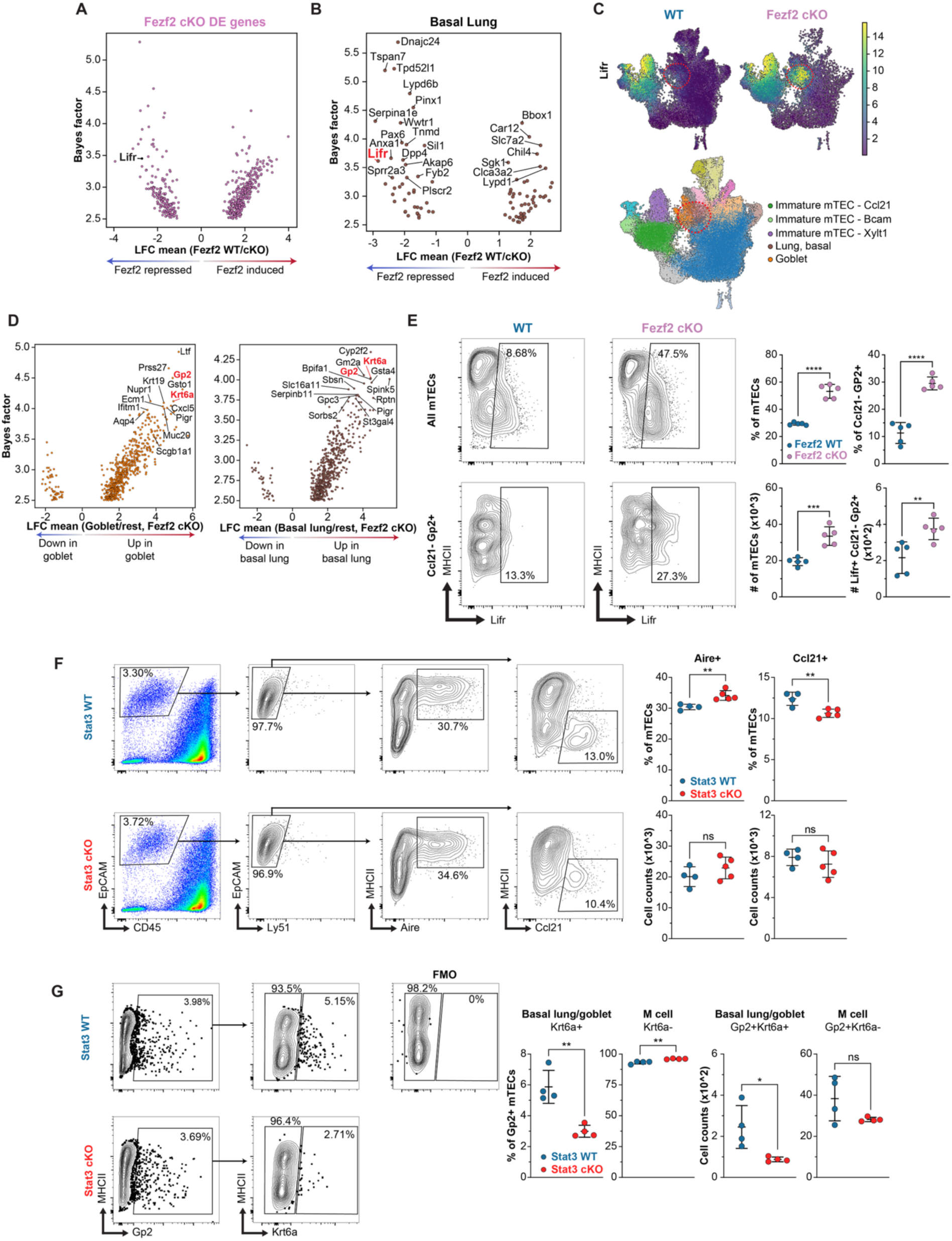
Lifr is transcriptionally repressed by Fezf2 in lung mimetic cells. (**A.**) Mean log fold change and bayes factors for Fezf2-regulated genes, highlighting Lifr. (**B.**) Mean log fold change and bayes factors for Fezf2-regulated genes in basal lung mimetic cells. (**C.**) Normalized expression of Lifr and its co-receptor in TECs. (**D.**) Highly differentially expressed genes for Fezf2 cKO goblet and basal lung mimetic cells vs all other mTECs. (**E.**) Representative flow cytometry gating of Lifr staining in WT and Fezf2 cKO mTECs (left) and quantification of the frequency and absolute number of all Lifr+ mTECs and Lifr+ Ccl21-Gp2+ mTECs across 6-week-old WT (n=5) and Fezf2 cKO (n=5) mice (right). (**F.**) Representative flow cytometry gating (left) and frequency or absolute count (right) of Aire and Ccl21-expressing mTECs in 6-week-old WT (n=4) and Stat3 cKO (n=5) mice. (**G.**) Representative flow cytometry gating (left) and frequency or absolute count (right) of basal lung/goblet (Gp2+Krt6a+) and microfold (Gp2+Krt6a-) mTECs in 6-week-old WT (n=4) and Stat3 cKO (n=5) mice. Statistical significance was calculated using unpaired Student’s t test (E-G).

Given that Lifr complex formation is upstream of Stat3 activation, we next wondered if disruption of TEC-intrinsic Stat3 signaling would display the converse of the Fezf2 cKO phenotype. Notably, previous work describing TEC development in Foxn1-Cre+/-Stat3fl/fl (Stat3 cKO) mice found a marked depletion of MHCII-low CD80-immature mTECs expressing Krt5 and Krt14 (32). To more precisely characterize these TEC alterations and align them to our evaluation of the Fezf2 cKO, we next utilized our spectral flow cytometry panel to enable quantification of both the immature (Ccl21+) and basal lung/goblet mimetic (Krt6a+ Gp2+) subsets. Consistently, Stat3 cKO mice had a decreased frequency of Ccl21+ mTECs, a decreased frequency and number of Gp2+ Krt6a+ mTECs, and a concomitant increase in the frequency of Aire+ mTECs (**Fig. 4F-G)**. Together, these findings are consistent with a mechanism in which Fezf2 antagonizes the Lif/Lifr/Stat3 signaling axis to modulate the steady-state balance between immature, Aire+ mTEC-hi, and lung epithelial mimetic subsets.

## DISCUSSION

Fezf2 is a Krüppel-type zinc finger protein whose only known functional regions are an N-terminal engrailed homology 1 motif (EH1 motif) followed by six C2H2-type zinc finger domains. EH1 motifs are distributed across many transcription factor domains and recruit transducin-like enhancer (Tle) family members. In turn, recruited Tles act as multidomain hubs for local transcriptional repression (33). Therefore, while there are reports of context-dependent direct transcriptional activation by Tles, the presence of an EH1 motif has typically been observed to mediate direct repression. In the central nervous system, Fezf2 has been intensively studied as a transcriptional switch guiding the molecular logic underlying corticothalamic subtype specification (25, 26). Here, the EH1 motif has been shown to recruit Tle4 to repress alternative neuronal programs (12). Given these results, a role for Fezf2 as a direct transcriptional repressor in mTECs is plausible. Previous efforts to identify Fezf2 binding partners in mTECs detected Chd4 by immunoprecipitation but, surprisingly, no Tle family members, despite their co-expression at the mRNA level in our single-cell RNA-seq datasets (7). Because commercially available antibody reagents to Fezf2 are limited, further interrogation of its binding partners and direct targets in mTECs may require improved tools. Finally, it is known that the Fezf2 promoter contains multiple conserved regulatory domains and that Sox4 and Sox11 are transactivators of Fezf2 expression in neurons (34). Sox4 is required for thymic tuft cell development and the expression of both Sox4 and Fezf2 are lymphotoxin-dependent in mTECs, suggesting parallel regulatory logic between neurons and mTECs that is partially downstream of Ltbr signaling in the thymus (6, 35).

An emerging theme in TEC biology is the definition of epithelial transcriptional regulatory networks underlying developmental access to mimetic subsets and interrogating their functional niches. Here we found that Fezf2 is required for the full program of mimetic cell development. While mimetic terminal differentiation serves to expand representation of mimetic-restricted TRAs, other functional contributions of mimetic subsets remain largely unknown. In our study, we found that iNKT2 polarization was disrupted in Fezf2 cKO thymus, consistent with diminished thymic tuft cell development and decreased IL-25 availability (23). A brief survey of thymic M cell biology revealed a Ccr6-dependent role in organizing the medullary B cell niche, reminiscent of peripheral Peyer’s Patches. In Fezf2 cKO mice, we found defects in B cell class switching, which are likely at least partially attributable to diminished M cell-derived Ccl20 and impaired B cell interaction with bona-fide thymic M cells. It is tempting to speculate that such spatially organized functional niches might also contribute to efficient negative selection. For example, it was recently shown that deletion of aquaporin 4 (Aqp4) reactive thymocytes requires B cell-intrinsic Aqp4 expression, despite substantial Aqp4 expression by TECs (36). Interrogation of whether B cell organization around thymic M cells alters the selection capacity of medullary B cells is an interesting area for further investigation.

Stat3 signaling during mTEC development has been implicated in the survival and maintenance of immature MHCII-low CD80-low mTECs (31, 37). This population is now understood to contain a large fraction of Ccl21+ immature TECs and may include progenitor subsets. We show that expression of Lifr, an IL-6 family cytokine receptor upstream of Stat3 activation, is highly enriched in Ccl21a-expressing immature mTECs and is transcriptionally repressed by Fezf2 in a subset of lung mimetic cells. Notably, both of these Lifr-expressing populations are significantly expanded in Fezf2 cKO mice. Together, these data suggest a mechanism by which Fezf2 modulates steady-state developmental dynamics and terminal differentiation of mTECs in part through Lifr repression and tuning of downstream Stat3 signaling tone. Therefore, definition of the role(s) of IL-6 family cytokine signaling and Jak/Stat pathway activation in TECs for stromal maintenance, as well as the sources of IL-6 family cytokines, are compelling areas for future work. However, while our model of direct Fezf2-mediated repression was able to rescue alterations in Ccl21+ and MHCII subset abundance, it was unable to normalize mimetic cell frequencies, and the possibility of direct transcriptional activation by Fezf2 remains incompletely resolved.

In summary, high-resolution single-cell transcriptomic analysis has revealed that Fezf2 partially functions as a transcriptional regulator of steady-state adult thymic epithelial development. While it shares this in common with Aire, the patterns of transcriptional control exerted are distinct. Both Aire and Fezf2 regulate the expression of TRAs in mTECs, but rather than driving a broad program of transcriptional induction, Fezf2 regulates a more limited program that also features prominent transcriptional repression. Mechanistically, Lifr was amongst the most repressed targets in aberrantly expanded lung mimetic cells and Stat3 cKO thymus displayed a converse phenotype to Fezf2 cKO, suggesting an unappreciated role for IL-6 family cytokine signaling in TEC developmental biology and an important area for future investigation.

## MATERIALS AND METHODS

### Study design

The objective of this study was to investigate mouse thymic epithelial cell biology using genetic murine model systems and high dimensional single-cell transcriptomics and integrative computational analyses.

### Mice

Aire-/-mice have been described previously (38). C57BL/6J (JAX 000664) and Stat3fl/f (JAX 016923) mice were purchased from the Jackson Laboratory. Foxn1-Cre and Fezf2bfl/fl mice were kindly provided by N.M (University of Georgia) and N.S. (Yale University) (39, 40). CD1 Fezf2 WT, total KO (tKO) and tKO-Engrailed transcriptional repressor domain (EnR) mice were generously shared and described by B.C. (University of California, Santa Cruz) and backcrossed for 4 generations to C57BL/6J (12). All mice were maintained in specific pathogen–free facilities at the University of California San Francisco (UCSF) in accordance with the guidelines established by the Institutional Committee on Animal Use and Care (IACUC) and Laboratory Animal Resource Center (LARC), and all animal procedures were approved by IACUC and LARC at UCSF. Age- and sex-matched mice between 3-12 weeks old were used for tissue harvest or experimental procedures unless otherwise specified in the text or figure legends.

### Single-cell tissue preparation

Mouse thymi were isolated, cleaned of fat and transferred to 10 ml of DMEM (Sigma-Aldrich D6546-6) with 2% Heat-Inactivated HyClone FBS (Fisher Scientific SH3040101) on ice. For mTEC and B cell enrichment, thymi were minced with a razor blade and tissue chunks were collected into 15-ml Falcon tubes, vortexed gently for 10 seconds and briefly spun down at 1250 RPM. Tissue fragments were resuspended in 4ml of DMEM digestion medium containing 2% FBS, 100ug/ml DNase I (Roche 10104159001) and 100ug/ml liberase TM (Sigma-Aldrich 5401127001). Tubes were incubated in a 37 °C water bath and tissue fragments were further digested with mechanical aid by pipetting through a glass Pasteur pipette every 4 min. At 8 min, tubes were spun briefly at 1250 RPM to pellet undigested fragments, and the supernatant was transferred to 20 ml of magnetic-activated cell sorting (MACS) buffer containing 0.5% BSA (Sigma-Aldrich), 2 mM EDTA (Teknova E0306-6) in PBS on ice to stop the enzymatic digestion. This procedure was repeated twice with 4 ml of fresh digestion medium each time, or until no visible tissue fragments remain. The single-cell suspension was then pelleted and washed once in MACS buffer and passed through a 70 μm filter. To enrich for mTEC and B cell subsets, density-gradient centrifugation using a three-layer Percoll gradient (Sigma-Aldrich E0414-1L) with specific gravities of 1.115, 1.065 and 1.0 was applied. Cells isolated from the Percoll-light fraction, between the 1.065 and 1.0 layers, were then resuspended in MACS buffer before counting and staining for flow cytometry.

### Flow cytometry and antibodies

Single-cell suspensions were prepared as described above and mixed with Live/Dead Fixable Blue Dead Cell Stain (ThermoFisher) in PBS for 20 min at 4°C followed by PBS wash and subsequent blocking with anti-mouse CD16/CD32 (24G2) (UCSF Hybridoma Core Facility) for 20 min at 4°C. Cells were then stained for surface markers with fluorochrome conjugated antibodies for 30 min at 4 °C in FACS buffer. For intracellular markers, cells were fixed and permeabilized with FoxP3 staining buffer kit (eBioscience) according to the manufacturer’s protocol. For intracellular Fezf2 RNA detection, PrimeFlow RNA assay kit (88-18005-204), mouse Fezf2 RNA paired probes and Alexa Fluor 568 label probes were purchased from Thermofisher Scientific and tested according to the manufacturer’s protocol. Flow cytometry data were collected on a LSRII Flow Cytometer (BD Biosciences) and Cytek Aurora (Cytek Biosciences) from the UCSF Single Cell Analysis Center and analyzed using FlowJo 10.8.1 software. The following antibodies were used in the study: AIRE (5H12), B220 (RA3-6B2), CCL21 (59106), CCR6 (29-2L17), CCR7 (4B12), CD4 (GK1.5), CD8 (53-6.7), CD11b (M1/70), CD11c (N418), CD19 (1D3), CD25 (3C7), CD45 (30-F11), CD69 (H1.2F3), CD73 (TY/11.8), CD80 (16-10A1), CD177 (FAB8186N), DCLK1 (EPR6085), EpCAM (G8.8), FoxP3 (MF-14), GP2 (2F11-C3), I-Ab (25-9-17), I-Ad (39-10-8), IgA (1040-09), IgD (11-26c.2a), IgG2b (RMG2b-1), IgM (RMM-1), Ly51 (6C3), MHC I (AF6-88.5.5.3), PLZF (Mags.21F7), Rorgt (Q31-378), SiglecF (1RNM44N), Tbet (4B10), TCRbeta (H57-597). Antibodies were purchased from Abcam, BioLegend, BD Biosciences, Novus Biologics, R&D Systems, Southern Biotech, MBL International, Fisher Scientific eBioscience, ThermoFisher Scientific or Miltenyi. For CD1d tetramer staining, mechanically dissociated thymocytes were incubated with BV421-conjugated PBS-57-loaded CD1d tetramer (NIH Tetramer Core) for 1 h at room temperature and washed twice with MACS buffer.

### Immunofluorescence staining and imaging

Mouse thymus was fixed in 2-4% paraformaldehyde (Thermo Scientific 28908) in PBS for 2 h at room temperature followed by overnight incubation at 4 °C in 30% (w/v) sucrose (Sigma-Aldrich S7903-1KG) in PBS. Tissues were embedded in Optimal Cutting Temperature Compound (Tissue-Tek 4583) and stored at −80 °C before sectioning (15–50 μm) on a cryostat (Leica).

Thin sections (15-50 μm) were mounted and dried on Superfrost Plus (Fisher Scientific 1255015) slides and moved directly to Immunomix solution containing 0.3% Triton X-100 (Sigma-Aldrich), 0.2% BSA (Sigma-Aldrich), 0.1% sodium azide (Sigma-Aldrich) in PBS. Thin section mounted slides were stained in a humidified chamber. Slides were rehydrated in PBS for 5 min before permeabilization in Immunomix and shaking for 1 h at room temperature followed by blocking with BlockAid (ThermoFisher, B10710), primary antibody staining at room temperature for 3 h, and, when needed, secondary antibody staining at room temperature for 1 h when sections were stained with unconjugated antibodies. DAPI staining was performed for 5 min at room temperature followed by 1X PBS wash for 5 min and repeated three times.

All sections were mounted with ProLong Diamond or Glass Antifade Mountant (ThermoFisher). Images were acquired on a Leica SP8 (Leica) laser scanning confocal microscope. The following primary antibodies were used in this study: B220 (RA3-6B2), Krt6a (polyclonal, cat # CL555-10590), and GP2 (2F11-C3). Antibodies were purchased from eBioscience, Life Technologies, and Proteintech. Secondary antibody: goat anti-rat AlexaFluor633 (A21094). Nuclear staining was performed with DAPI (Biolegend 422801).

### Single-cell RNA sequencing data processing

Raw sequencing reads (.fastq files) from four WT, two Aire KO and two Fezf2 cKO samples were aligned using kallisto-bustools (v0.27.3) to the prebuilt ensembl v96 mouse reference genome for kallisto with the default parameters (41). The resulting per-cell, per-gene UMI counts were further processed in Python (v3.10.13) using scanpy (v1.9.3) for all downstream analysis (42). Transcripts were first filtered on genes with protein coding biomart annotations followed by a preliminary filter step removing all cells with less than 100 genes and all genes detected in fewer than 20 cells. Scrublet (v0.2.3) was then used to remove cells likely to be doublets or multiplets by excluding cells with doublet scores over 0.2 from the data (43). Samples were further filtered to exclude low quality or damaged cells with over 5% mitochondrial reads, more than 70,000 or less than 1,000 UMIs, and over 8,500 or less than 1,000 genes detected. After filtering and merging all samples, 53,540 cells remained for downstream analysis along with 19,866 genes. The python package scvi-tools (v1.0.4) was used to train an scVI model of gene expression for these genes across all cells to normalize raw counts data and integrate each sample into a single dataset with batch correction (17). Batch correction performance was quantified on each genotype using the scib (v1.1.4) package (44). The scVI latent space was used to identify nearest neighbors and perform dimensional reduction for data visualization using scanpy’s umap function. Normalized read counts for each cell were calculated using scvi-tools’ get_normalized_expression function with library_size=10,000 UMIs per cell. Leiden clustering was performed using scanpy with resolution=0.4 and three clusters containing non-mTEC cell populations were removed from the dataset leaving 50,340 cells. After recomputing the UMAP for the remaining mTEC cells, cell types were annotated using celltypist (v1.6.2) label transfer from a published mimetic cell scRNA-seq dataset and refined by subclustering (9, 18). Scvi-tools normalized counts were used for all downstream feature plot visualizations unless otherwise specified. UMAP feature plots were generated by clipping the minimum and maximum colormap values for gene expression to the 1st and 99th percentile of scVI normalized expression for that gene to prevent outlier gene expression from influencing visualizations.

### Cell type annotation with celltypist

A published scRNA-seq dataset enriched for post-Aire mimetic cells with high resolution cell type annotations was used for reference label transfer in our analyses (9). Raw count data from the refence dataset were filtered on all annotated cells and transcripts with protein coding biomart annotations that were detected in at least 20 cells. Raw data was library size scaled to 10,000 counts per cell and log-transformed with the scanpy “normalize_total” and “log1p” methods, respectively. A celltypist (v1.6.2) label transfer model was trained on the top 4480 genes using the “train” method with default parameters and 1000 maximum iterations (18). This label transfer model was then used to annotate cell types in our analyses with similarly library size scaled and log-transformed raw data using the celltypist “annotate” method with majority_voting set to True. Immature MEC, Aire-stage, and TA mTEC cells were separately subclustered using leiden clustering with resolution 0.2, 0.2, and 0.1, respectively. Highly differentially expressed genes for each subcluster vs all other WT mTECs were identified with scVI’s differential_expression method as genes with bayes factor greater than 3, predicted mean log fold change greater than 1 or less than -1, and non-zero expression in at least 10% of the subcluster of interest. Immature mTECs were then split into 3 new subclusters defined by their uniquely high expression of Bcam, Ccl21, and Xylt1 in this differential gene expression analysis (**Fig S1E-F**). Other subclusters of the immature and Aire-stage mTECs misannotated by celltypist were relabeled as existing TEC subsets based on marker gene expression (Dclk1, Krt15, Foxj1, Aire, and Prss16).

### Differential abundance of WT and KO mTEC subsets

The abundance of each cell type in samples for each of our datasets was determined by taking the fraction of all cells in each sample that are assigned that label in that sample. The relative density of WT and KO cells in UMAP latent space was determined using the kernel density of the UMAP latent representation of each sample as previously described (45). In short, this was calculated using the statsmodels (v0.14.0) KDEMultivariateConditional kernel density estimator with the UMAP coordinates for each cell as continuous dependent variables, the genotype of each cell as an unordered independent variable and the bandwidth set to the normal_reference default. The log ratio of resulting estimated WT and KO densities for each point in the UMAP latent space takes on positive values for cells that fall into regions of the UMAP latent space that have a higher density of KO cells, negative values for cells that fall into regions that have a higher density of WT cells, and values close to 0 where both samples are equally dense.

### Differential expression

For WT vs KO comparisons, differential expression analysis was performed separately for each cell type with at least 30 cells in each Aire KO and Fezf2 cKO sample relative to the WT control samples to identify Aire and Fezf2 regulated genes across all datasets. The scvi-tools differential_expression function was used with the scVI model trained for the merged dataset to calculate Bayes factors describing how likely each gene in the dataset is to be differentially expressed between the WT and each KO sample (17). Highly differentially expressed genes for a given cell type were classified as genes with a Bayes factor over 2.5, a predicted mean log2 fold change of expression greater than 1 or less than -1 compared to the WT sample, at least 5% of cells with nonzero expression, and peak (99th percentile) normalized expression in that cell type over 0.5. The same approach was used to identify differentially expressed genes between individual WT and between individual KO replicates to filter out genes that were attributable to batch effects. Additionally, gene/cell type combinations with opposite effects across samples (i.e. Fezf2-induced in sample 1 but Fezf2-repressed in the same cell type in sample 2) were removed. The final lists of Fezf2- and Aire-regulated genes were generated by taking the intersection of differentially expressed genes for each WT vs KO replicate comparison and removing any genes that were differentially expressed in the corresponding WT vs WT and KO vs KO comparisons. For volcano plots, if a gene was differentially expressed in more than one mTEC subset, the highest log2 fold change and corresponding bayes factor was used. For goblet and basal lung vs all other mTECs comparisons in the Fezf2 cKO samples, highly differentially expressed genes were identified as genes with a bayes factor greater than 2.5, a mean log fold change greater than 1 or less than -1, and expression in at least 10% of the population of interest.

### Defining tissue restricted antigens

The cellxgene mouse tissue atlas was filtered on transcripts with protein coding biomart annotations that were detected in at least 20 cells as well as cells with at least 500 detected genes. An scVI model of gene expression was generated using the original batches annotated for each dataset in the atlas. Normalized expression was reconstructed using the scVI model with a library size of 10,000 counts. For TRA scoring, genes detected with at least 2 raw counts in at least 100 cells were retained. Normalized expression of the detected genes was averaged across tissue or cell type annotations, squared, and transformed to a probability such that expression for each gene across all tissues or cell types sums to 1. The Shannon entropy or Tau score of the resulting probability distribution was then calculated as previously described (46, 47). Finally, the Shannon entropy of each gene (H(g)) was converted to a score ranging from 0 (equal expression in all tissues/cell types) to 1 (expressed only in a single tissue/cell type) using the equation:

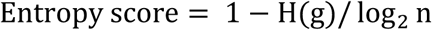

where n is the number of tissues/cell types in the cellxgene atlas. To classify genes as TRAs or non-TRAs, a logarithmic model was fit to the Shannon entropy score and Tau score relationship for each gene. This model was used to determine the appropriate Shannon entropy score threshold corresponding to each chosen Tau score threshold (0.85, 0.9, and 0.95) before both thresholds were used to define TRAs and non-TRAs for each scoring metric (**Fig. S3A**). Scoring using gene expression across cell types identified additional genes with restricted expression that were not classified as TRAs using tissue expression, so cell type TRAs were used for all analyses in this study (**Fig. S3B**). Shannon entropy and Tau cell type TRA scores were consistent with a previously published TRA list (**Fig. S3C**) (48). For our analyses, a gene was classified as a TRA if it passed either the Tau or Shannon entropy score threshold as genes that were called TRAs by only one method were still largely restricted in their expression patterns across cell types (**Fig. S3D**).

### Statistical analysis

All statistical analysis was performed using Prism 9 (GraphPad Software) unless stated otherwise. Statistical significance between two groups was calculated using an unpaired, parametric, two-tailed Student’s t test. If more than two groups were plotted together, statistical significance was calculated either by using two-way non-parametric ANOVA (Sidak’s test) with multiple comparisons. Experimental groups included a minimum of two to three biological replicates. Intragroup variation was not assessed. Figures display mean ± s.d. unless otherwise noted. A p value of less than 0.05 was considered statistically significant with *p< 0.05, **p<0.01, ***p<0.001, and ****p<0.0001. Specific statistical tests for each figure used are referenced in figure legends. No statistical methods were used to predetermine sample size.

## Data Availability

All new scRNA-seq sequencing datasets generated for analysis have been uploaded to GEO under the accession number GSE226493. Existing public datasets used for this study include scRNA-seq of post-Aire mimetic cell enriched mTECs (GSE194253), non-T thymic hematopoietic cells from Dex-treated mice (GSE244229), and the cellxgene mouse tissue atlas (9, 30, 49).

## Code Availability

All code files used to process data and generate figures for this study are available at https://github.com/j-germino/fezf2-thymus.git.

## Funding

NIH R01MH094589 and NIH R01NS089777 (BC)

ARCS Foundation, Northern California (JG)

National Institute of Allergy and Infectious Disease R37AI097457 (MSA) UCSF Sandler Fellows PSSP grant (JMG)

UCSF T32 Translational Science (XL)

National Institute of Aging grant K01AG072789 (CNM)

W.M. Keck Foundation grant (JMG)

Pew Biomedical Scholars Program (JMG)

## Author contributions

Conceptualization: MSA, JMG, CNM

Formal Analysis (bioinformatics): JG, VN

Methodology: CNM, JMG, VN, BC, SV, YW, XL, JG

Investigation: XL, JG, YW, JB, DY, YN, JT

Visualization: XL, JG

Funding acquisition: MSA, JMG

Project administration: IP

Supervision: VN, MSA, JMG, CNM

Writing – original draft: XL, JG, JMG, CNM

Writing – review & editing: XL, JG, VN, MSA, KR, BC, JMG, CNM

## Supplementary Information

### Supplementary Tables

**Table S1. Immature mTEC subset highly differentially expressed genes in WT mTECs.** Highly differentially expressed genes for each immature mTEC subcluster in WT mTECs were calculated using scvi-tools, identified as genes with a bayes factor greater than 3, a mean log-fold change between the population of interest and all other WT mTECs greater than 1 or less than -1, and expression in at least 10% of the immature mTEC subcluster.

**Table S2.**
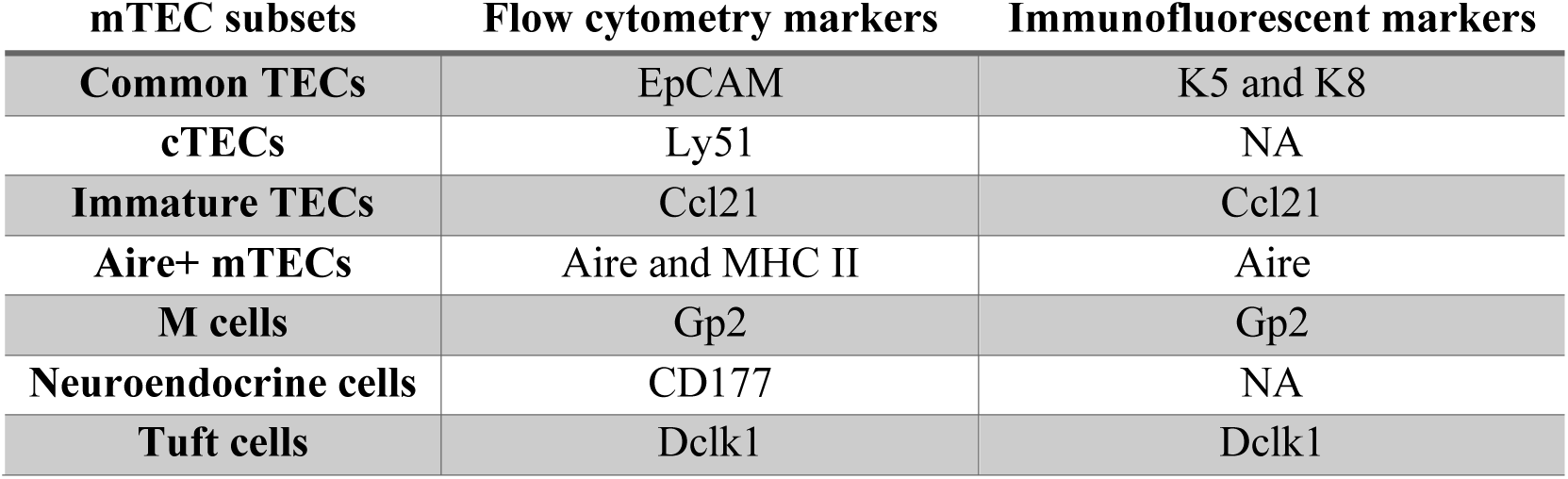
Flow cytometry marker genes used for mTEC subpopulation identification.

| mTEC subsets | Flow cytometry markers | Immunofluorescent markers |
| --- | --- | --- |
| **Common TECs** | EpCAM | K5 and K8 |
| **cTECs** | Ly51 | NA |
| **Immature TECs** | Ccl21 | Ccl21 |
| **Aire+ mTECs** | Aire and MHC II | Aire |
| **M cells** | Gp2 | Gp2 |
| **Neuroendocrine cells** | CD177 | NA |
| **Tuft cells** | Dclk1 | Dclk1 |

**Table S2. Flow cytometry marker genes used for mTEC subpopulation identification.**

**Table S3. WT vs Aire KO highly differentially expressed genes.** Highly differentially expressed genes between WT and Aire KO samples for each cell type were calculated using scvi-tools, identified as genes with a bayes factor greater than 2.5, a mean log-fold change between WT and Aire KO cells greater than 1 or less than -1, and expression in at least 5% of either sample while also excluding genes with low maximum expression. differentially expressed genes were filtered on genes that were consistent across both Aire KO replicates and not differentially expressed between WT or KO samples in any cell type.

**Table S4. WT vs Fezf2 cKO highly differentially expressed genes.** Highly differentially expressed genes between WT and Fezf2 cKO samples for each cell type were calculated using scvi-tools, identified as genes with a bayes factor greater than 2.5, a mean log-fold change between WT and Fezf2 cKO cells greater than 1 or less than -1, and expression in at least 5% of either sample while also excluding genes with low maximum expression. differentially expressed genes were filtered on genes that were consistent across both Fezf2 cKO replicates and not differentially expressed between WT or KO samples in any cell type.

**Table S5. TRA analysis of genes in the cellxgene scRNA-seq atlas.** The expression of each gene in the cellxgene scRNA-seq atlas was averaged per cell type, and the resulting mean expression distribution across cell types was used to compute Tau and Shannon entropy scores per gene for TRA classification (**see methods**).

### Supplementary Figures

**Figure S1.**
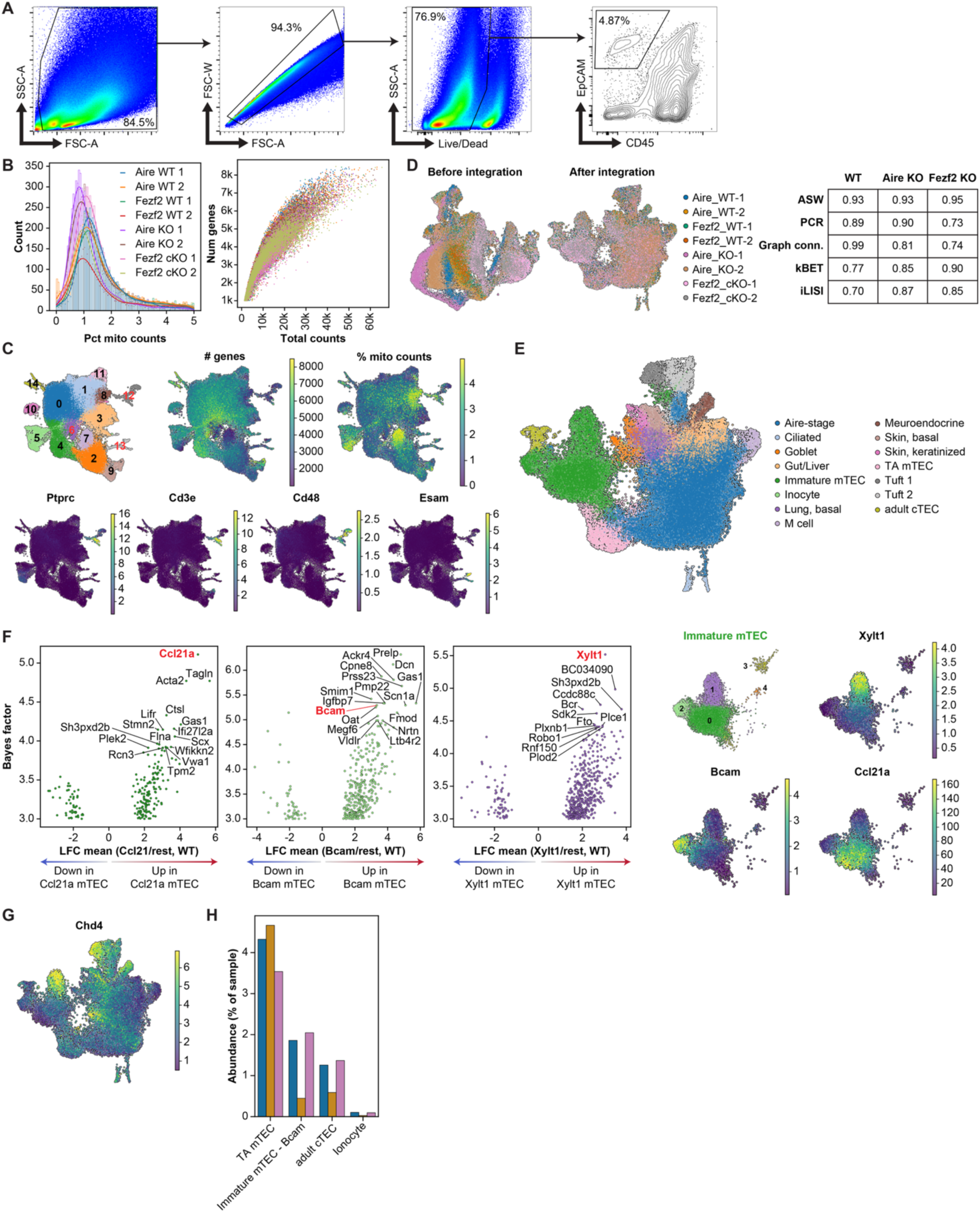
Collection and annotation of WT, Fezf2 cKO and Aire KO single-cell RNA-seq datasets. (**A.**) Sorting strategy of EpCAM+ CD45-TECs for 10X scRNA-seq. (**B.**) Per-cell mitochondrial read percentage (left) and total counts versus number of genes detected (right) colored by batch for 4-6-week-old WT, Aire KO, and Fezf2 cKO EpCAM+ CD45-TECs scRNA-seq datasets. (**C.**) Feature plots depicting the cluster number, number of detected genes, percent mitochondrial reads, and normalized expression for non-TEC markers. Cluster numbers in red were non-TECs removed from downstream analysis. (**D.**) Feature plots colored by sample before and after batch correction. Table depicts integration QC metrics calculated using scib scaled to range from 0 (poor integration) to 1 (well-integrated) (44). (**E.**) Predicted cell type labels of individual cells determined using celltypist reference label transfer from a published mTEC scRNA-seq dataset sampling mimetic cells with high resolutions (9). (**F.**) Differentially expressed genes analysis (left) and feature plots of normalized expression for marker genes (right) used to identify and reannotate immature mTEC subclusters. (**G.**) Normalized expression of Chd4 in WT TECs. (**H.**) Fraction of cells annotated as each TEC subpopulation across the WT, Aire KO, and Fezf2 cKO samples for subpopulations with a mean log2(fold change) between WT and Fezf2 cKO samples less than 0.5 or greater than -0.5. Colored bar depicts the mean abundances across 2 (Fezf2 cKO and Aire KO) or 4 (WT) independent replicates.

**Figure S2.**
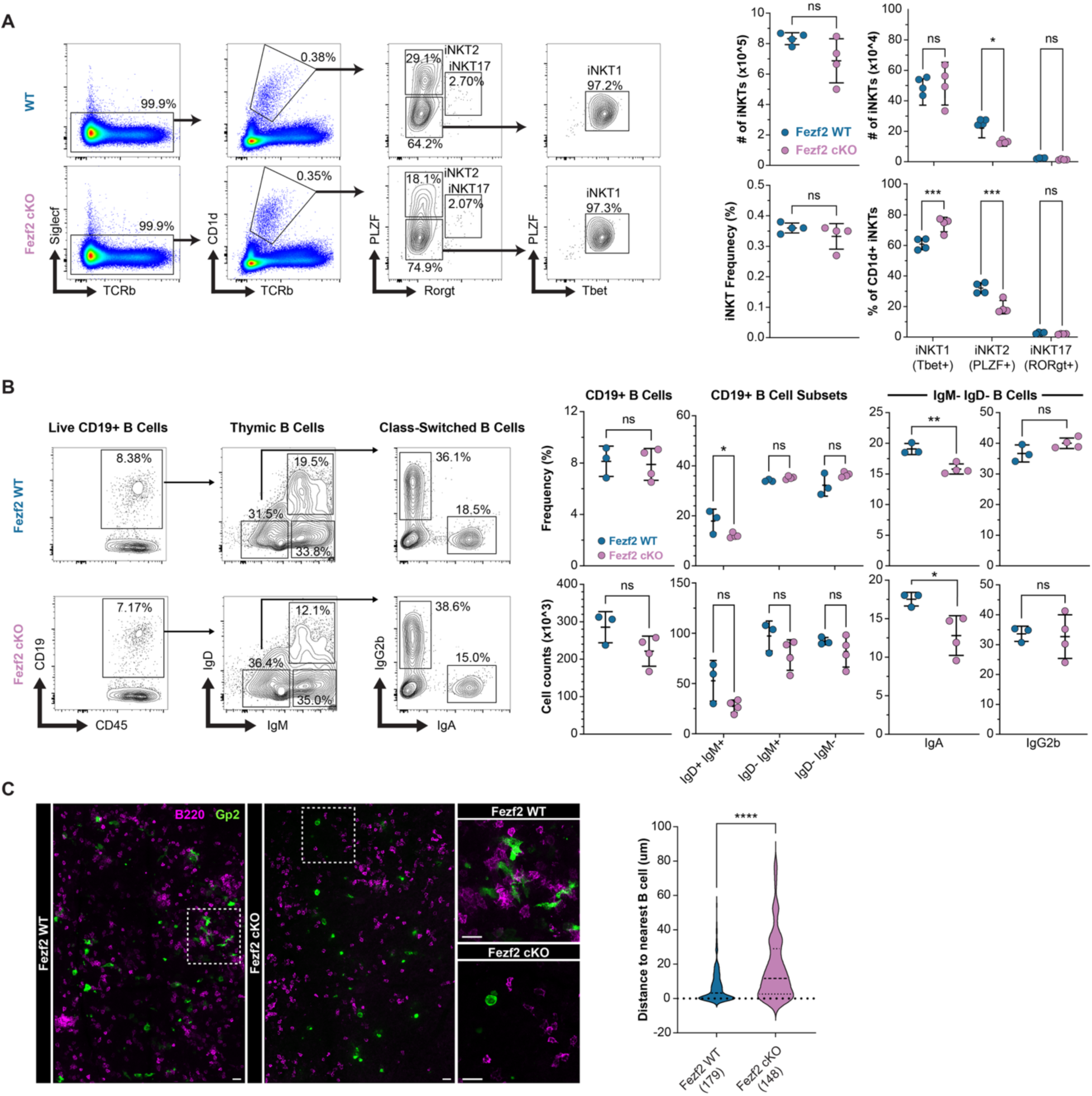
Developmental defects in Fezf2 cKO thymus disrupt stromal-immune crosstalk. (**A.**) Representative FACS plots of each population (left) along with frequency or absolute number of iNKTs (middle) and iNKT subsets (right) in 6-week-old WT (n=4) and Fezf2 cKO (n=4) mice. (**B.**) Representative FACS plots (left) and frequency or total counts (right) of thymic B cell subsets in 6-week-old WT (n=4) and Fezf2 cKO (n=4) mice. (**C.**) Representative immunofluorescent staining of Gp2 and B220 in 6-week-old WT and Fezf2 cKO thymi (left) and quantification of the distance between Gp2+ mTECs and the nearest B cell (right) (n=3). Scale bars are 20 um (C), and 10 um (C, inserts). Statistical significance was calculated using unpaired Student’s t test (A-B).

**Figure S3.**
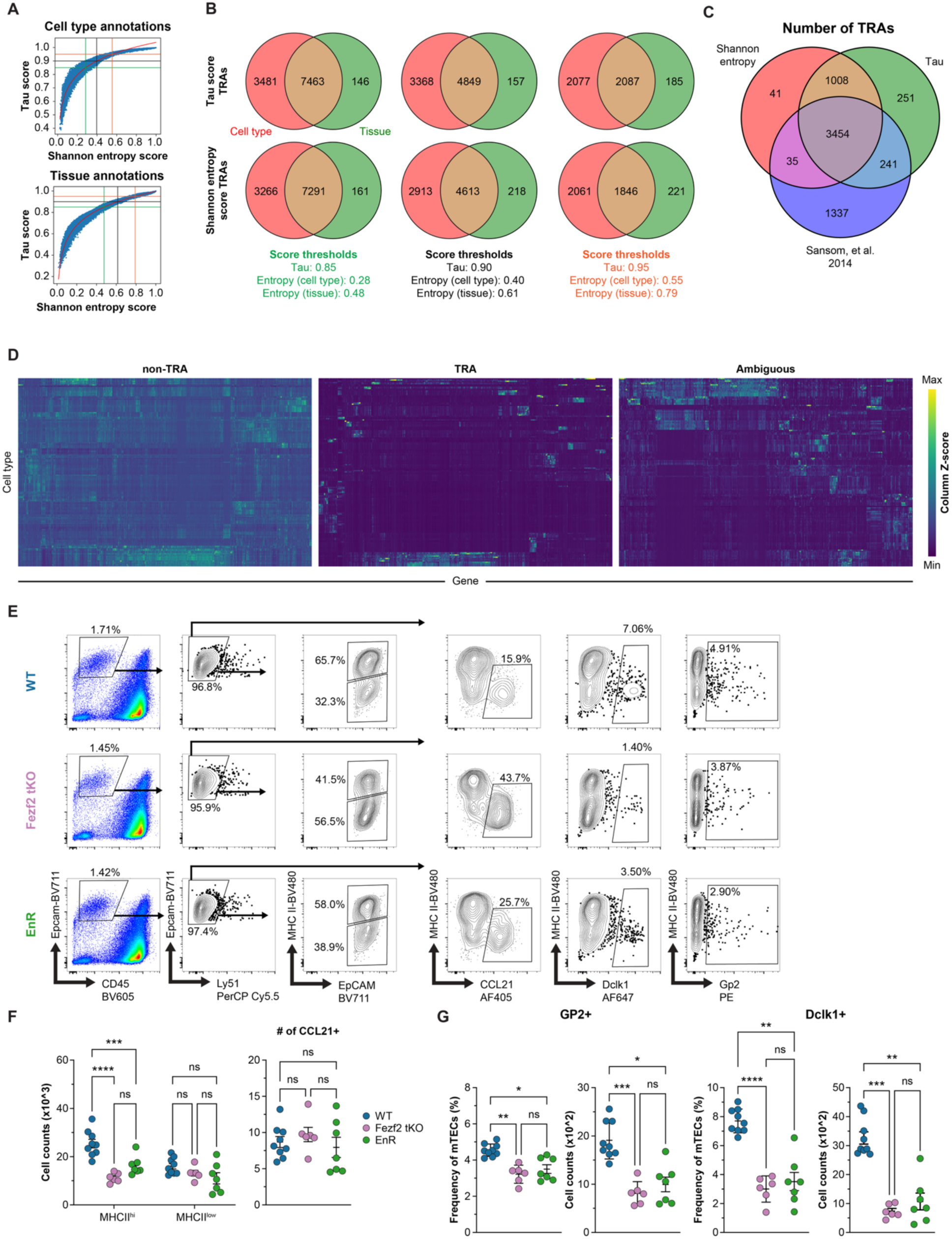
Identifying TRAs using the cellxgene scRNA-seq atlas. (**A.**) Shannon entropy vs Tau scores generated from cell type and tissue annotations in the cellxgene atlas (49). Vertical and horizontal lines depict score thresholds used for TRA analyses. Red line shows a log2 model fit to determine the appropriate Shannon entropy score for each Tau threshold. (**B.**) Overlap in TRA gene lists between cell type- and tissue-level cellxgene atlas scoring. (**C.**) Overlap between TRAs from Tau and Shannon entropy scoring of cellxgene cell types and a previously published TRA list (48). (**D.**) Cell type expression in the cellxgene atlas of TRAs/non-TRAs identified by both scoring metrics (left, middle) or by one scoring metric (right). (**E.**) Representative flow cytometry gating strategy for mTEC subpopulations across WT, Fezf2 tKO, and Fezf2 tKO EnR mice. (**F.**) Absolute number of major mTEC subpopulations across 4- to 7-week-old WT, Fezf2 tKO and Fezf2 tKO EnR mice. (n=6-9 mice). (**G.**) Frequency and absolute number of mimetic cell populations across 4- to 7-week-old WT, Fezf2 tKO and Fezf2 tKO EnR mice. (n=6-9 mice). Statistical significance was calculated using two-way ANOVA (Sidak’s test) with multiple comparisons (F-G).

**Figure S4.**
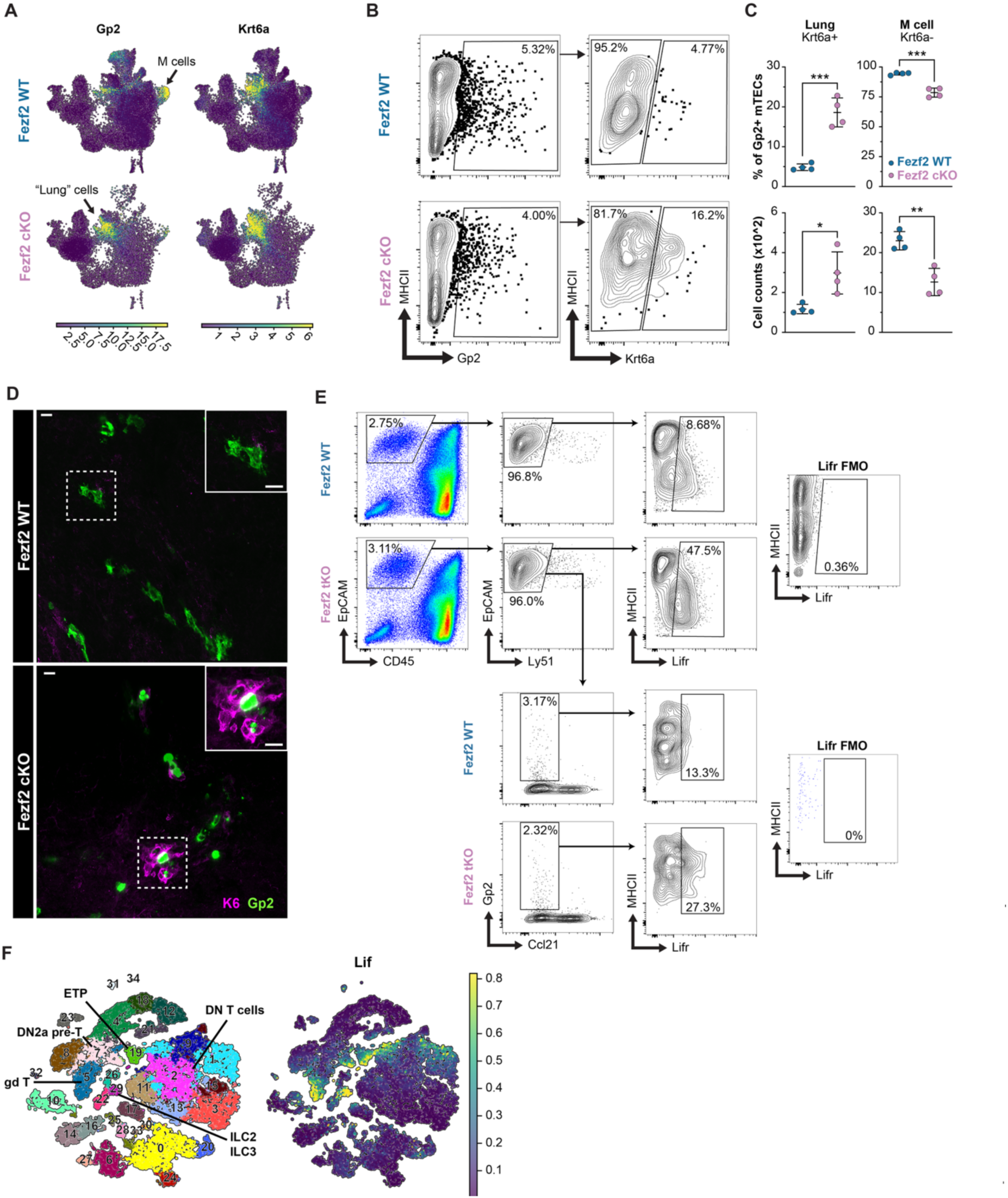
Lifr expression in the murine thymus. (**A.**) Feature plots of normalized Gp2 and Krt6a expression in WT and Fezf2 cKO TECs. (**B.**) Representative flow cytometry gating for basal lung/goblet (Krt6a+Gp2+) and microfold (Krt6a-Gp2+) mTECs. (**C.**) Quantification of frequency and count of basal lung/goblet and microfold mimetic cells in 6-week-old WT (n=4) and Fezf2 cKO (n=4) thymi by flow cytometry. (**D.**) Immunofluorescent staining of Gp2 and Krt6a in WT and Fezf2 cKO mice. Scale bars are 50um. (**E.**) Representative flow cytometry gating used to identify Lifr-expressing mTECs. (**F.**) Lif expression in hematopoietic cells from the murine thymus before and after Dex treatment (31).

